# Chromosomal-level genome of the *Pinus massoniana* ‘Minlin WY36’

**DOI:** 10.1101/2025.06.23.660646

**Authors:** Wei-Hong Sun, Feiping Zhang, Xuewei Zhao, Zhen Li, Ke-Wei Liu, Songqing Wu, Yajie Guo, Xia Hu, Mengyao Zeng, Linying Wang, Cheng-Yuan Zhou, Deqiang Chen, Ruiyue Zheng, Xiaopei Wu, Dong-Hui Peng, Liang Ma, Zhi-Wen Wang, Wenfeng Lan, Huimin Chen, Shuangquan Zou, Wen-Chieh Tsai, Yves Van de Peer, Zhong-Jian Liu, Siren Lan

## Abstract

Owing to conifers complex and large genomes, it is challenging to construct a comprehensive reference genome and conduct evolutionary and genomic research. Here, we present the chromosome-level assembly of *Pinus massoniana* ‘Minlin WY36’, a pioneering conifer species used for afforestation, timber production, and oleoresin extraction. The genome size was determined to be 23.41 Gb, with a contig N50 value of 1.15 Mb and a scaffold N50 value of 2.02 Gb. A total of 23.16 Gb of reads (98.93% of the assembled genome) were anchored and oriented onto 12 pseudochromosomes, with chromosome lengths from 1.47 to 2.29 Gb. A total of 32,144 protein-coding genes were predicted, of which 27,173 (84.54%) were functionally annotated. The *P. massoniana* ‘Minlin WY36’ genome consists of 79.41% (18.57 Gb) repetitive sequences, among which long terminal repeat sequences (LTR) were the most abundant. Through comparative genomic analysis, we found that the unique and significantly expanded genes in *P. massoniana* ‘Minlin WY36’ were enriched in the metabolism related to resin synthesis. This study offers valuable insights into conifer evolution and contributes resources for further investigations of conifer adaptation and developmental processes.

## Introduction

Pines (genus *Pinus*) belong to the coniferous family Pinaceae of gymnosperms, with approximately 110–120 species distributed across the Northern Hemisphere (Gernandt *et al*., 1999). Among these, *Pinus massoniana* is a unique conifer species endemic to China (He *et al*., 2023). Its planting area ranks first in the distribution of tree species in China, and is widely distributed in southern China. As a crucial species for afforestation, timber production, and oleoresin extraction, *P. massoniana* has extensive applications in construction, papermaking, and pharmaceutical industries and has important ecological and economic significance (Peng *et al*. 2003; Zhang *et al*. 2013). In this study, we generated a high-quality *P. massoniana* ‘Minlin WY36’ genome sequence to elucidate the evolution of its genome structure and phylogenetic relationships within gymnosperms. This study offers profound insights into the evolution of conifers and serves as a foundation for the further exploration of their adaptive developmental strategies.

## Results and discussion

Survey analysis revealed a high level of heterozygosity in the *P. massoniana* ‘Minlin WY36’ genome, accounting for 1.13% of the 23.38 Gb genome size determined by *K*-mer analysis. For *de novo* whole-genome sequencing of *P. massoniana* ‘Minlin WY36’, we obtained 2165.08 Gb of clean reads with an average length of 12.47 Kb using PacBio technology. The final assembled genome amounted to 25.37 Gb with a contig N50 value of 1.15 Mb (**Table 1**). High-throughput and high-resolution chromosome conformation capture (Hi-C) technology was used to obtain a chromosome-level genome. Based on the Hi-C data, the genome was assembled to a total size of 23.41 Gb with a scaffold N50 value of 2.02 Gb (**Table 1**). A total of 23.16 Gb reads were mapped to 12 chromosomes, with the chromosome lengths ranging from 1.47 to 2.29 Gb (**Table 2**).

**Table 1.**
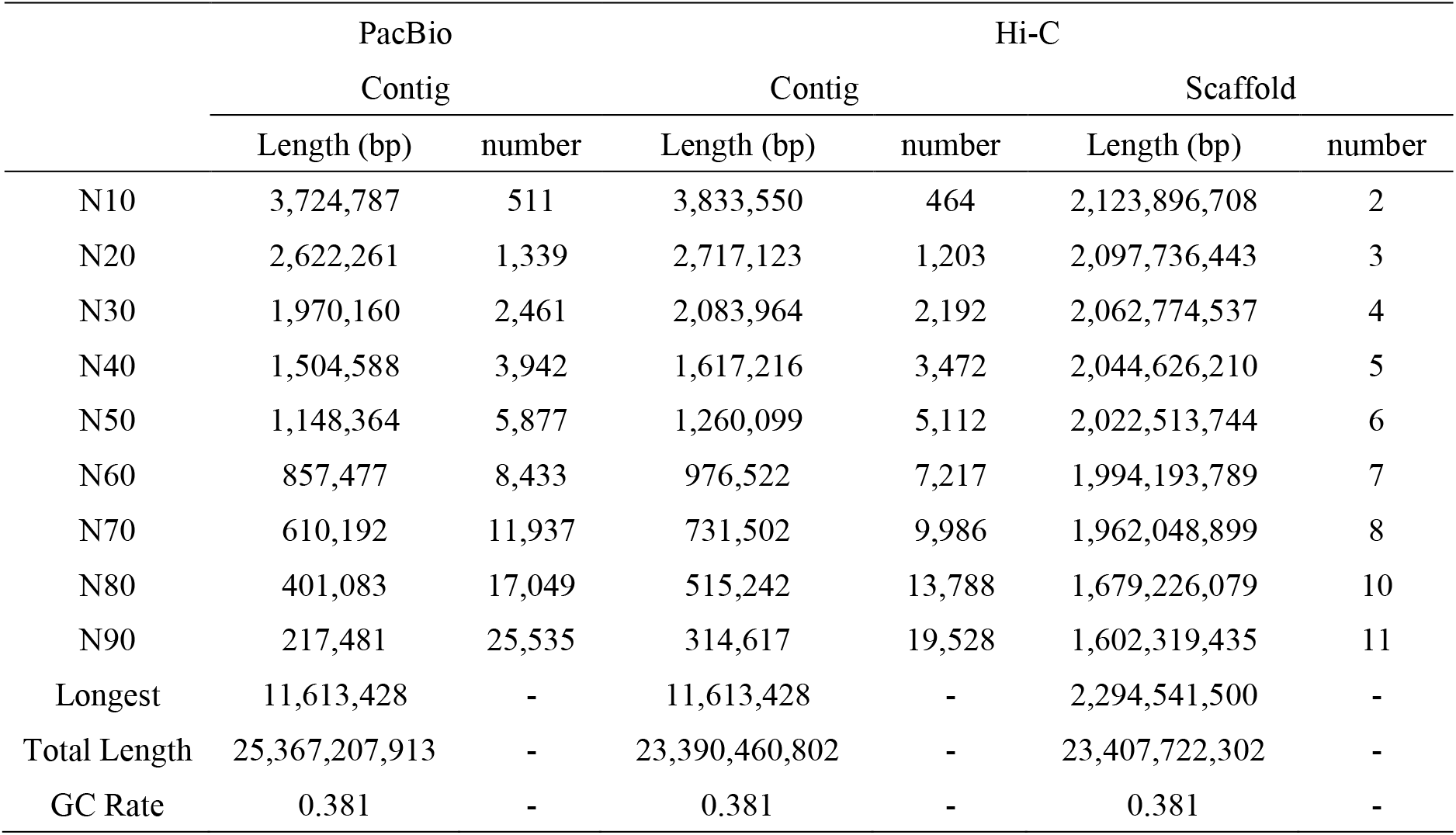
The statistic of genome assembly of *P. massoniana* ‘Minlin WY36’.

|  | PacBio |  |  | Hi-C |  |  |
| --- | --- | --- | --- | --- | --- | --- |
|  | Contig |  | Contig |  | Scaffold |  |
|  | Length (bp) | number | Length (bp) | number | Length (bp) | number |
| N10 | 3,724,787 | 511 | 3,833,550 | 464 | 2,123,896,708 | 2 |
| N20 | 2,622,261 | 1,339 | 2,717,123 | 1,203 | 2,097,736,443 | 3 |
| N30 | 1,970,160 | 2,461 | 2,083,964 | 2,192 | 2,062,774,537 | 4 |
| N40 | 1,504,588 | 3,942 | 1,617,216 | 3,472 | 2,044,626,210 | 5 |
| N50 | 1,148,364 | 5,877 | 1,260,099 | 5,112 | 2,022,513,744 | 6 |
| N60 | 857,477 | 8,433 | 976,522 | 7,217 | 1,994,193,789 | 7 |
| N70 | 610,192 | 11,937 | 731,502 | 9,986 | 1,962,048,899 | 8 |
| N80 | 401,083 | 17,049 | 515,242 | 13,788 | 1,679,226,079 | 10 |
| N90 | 217,481 | 25,535 | 314,617 | 19,528 | 1,602,319,435 | 11 |
| Longest | 11,613,428 | - | 11,613,428 | - | 2,294,541,500 | - |
| Total Length | 25,367,207,913 | - | 23,390,460,802 | - | 23,407,722,302 | - |
| GC Rate | 0.381 | - | 0.381 | - | 0.381 | - |

**Table 2.**
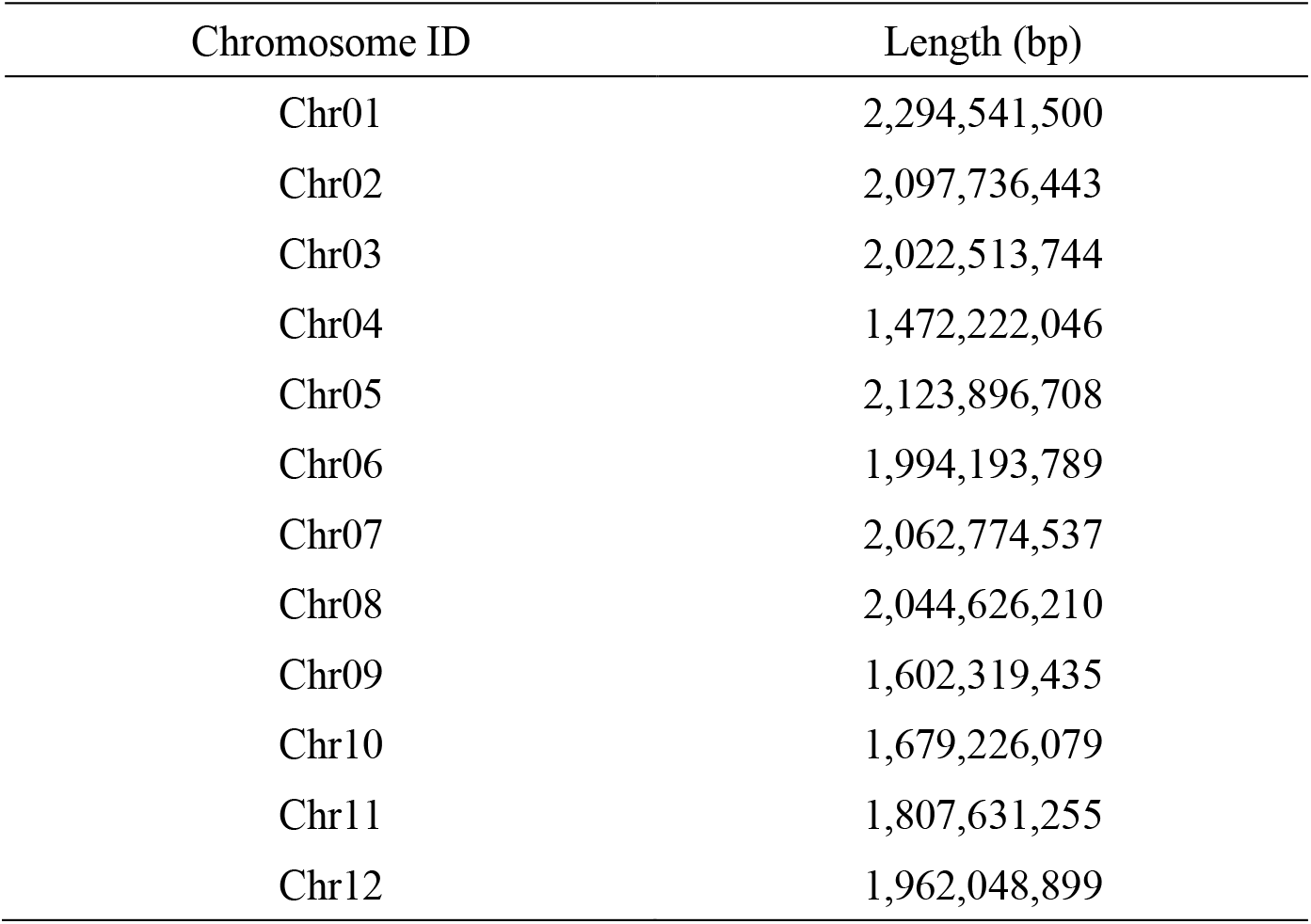
The length of chromosome by Hi-C assemble of *P. massoniana* ‘Minlin WY36’.

| Chromosome ID | Length (bp) |
| --- | --- |
| Chr01 | 2,294,541,500 |
| Chr02 | 2,097,736,443 |
| Chr03 | 2,022,513,744 |
| Chr04 | 1,472,222,046 |
| Chr05 | 2,123,896,708 |
| Chr06 | 1,994,193,789 |
| Chr07 | 2,062,774,537 |
| Chr08 | 2,044,626,210 |
| Chr09 | 1,602,319,435 |
| Chr10 | 1,679,226,079 |
| Chr11 | 1,807,631,255 |
| Chr12 | 1,962,048,899 |

To predict gene models, we used a combination of *de novo* prediction, homology-based searches, and RNA-sequencing data, resulting in the prediction of 32,144 protein-coding genes in *P. massoniana* ‘Minlin WY36’ genome. The primary gene models exhibited a mean length of 37,832.39 bp, with an average exon length of 263.80 bp and an average intron length of 12,804.88 bp. Similar to other gymnosperms with large genomes, extremely long introns were also predicted in *P. massoniana* ‘Minlin WY36’ (**Figure 1A**). This indicates that the accumulation of long introns has led to a significant expansion of the genome (Nystedt *et al*., 2013). On the basis of a Benchmarking Universal Single-Copy Orthologs (BUSCO) assessment, the completeness of the gene annotations was approximately 81.75%. We functionally annotated the protein-coding genes using various databases, including Gene Ontology (GO), Kyoto Encyclopedia of Genes and Genomes (KEGG), Swiss-Prot, and others, an over 84.54% of the genes could be mapped to corresponding functional items. These analyses indicate a relatively comprehensive structural annotation of the reference genome. A total of 113 microRNAs (miRNAs), 730 small nuclear RNA (snRNAs), 2,011 transfer RNAs (tRNAs), and 3,307 ribosomal RNAs (rRNA) were identified in the *P. massoniana* ‘Minlin WY36’ genome (**Table 3**). We estimated that 79.41% (18.57 Gb) of the *P. massoniana* genome was composed of repetitive sequences, among which transposable elements (TE) sequences were relatively abundant, with a long terminal repeat (LTR) accounting for 70.94%, followed by LINE accounting for 4.88%, and DNA accounting for 2.34%. We used complete LTR-Copia and LTR-Gypsy elements from *P. massoniana* ‘Minlin WY36’ to investigate further the timing of conifer transposable element insertions and found that insertions appear to have occurred tens of millions of years ago (**Figure 1B**).

**Figure 1.**
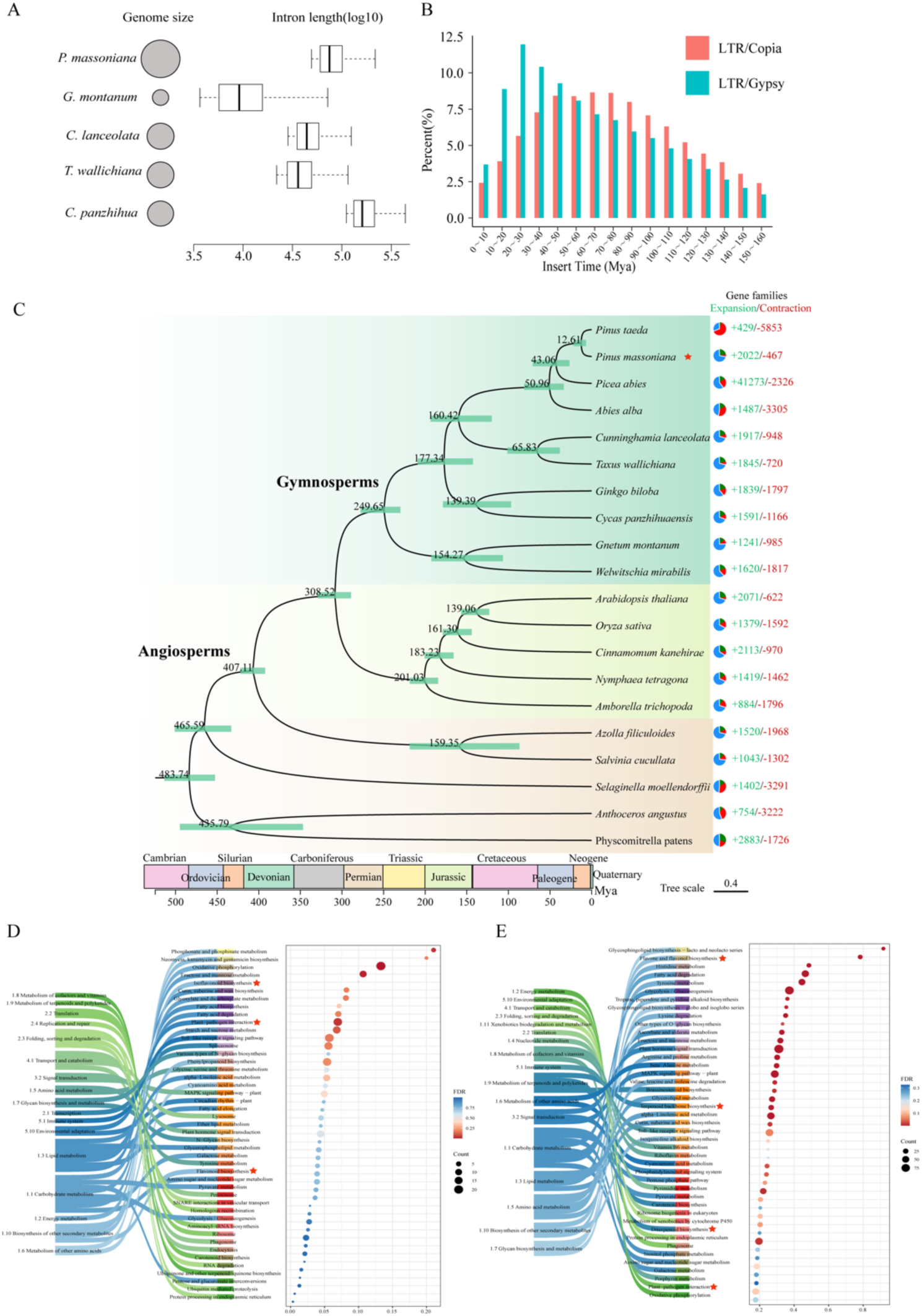
Assembly and genome features of *Pinus massoniana*. **A**. Intron length. **B**. Long terminal repeat insertion time in *P. massoniana* ‘Minlin WY36’. **C**. Phylogenetic tree showing divergence times and evolution of the gene family size in the 20 species. Green and red numbers indicate the number of expanded and contracted gene families, respectively. The blue portions of the pie charts represent gene families with constant copy number. **D**. KEGG enrichment of genes unique to *P. massoniana* ‘Minlin WY36’. **E**. KEGG enrichment of significantly expanded genes in *P. massoniana* ‘Minlin WY36’.

**Table 3.**
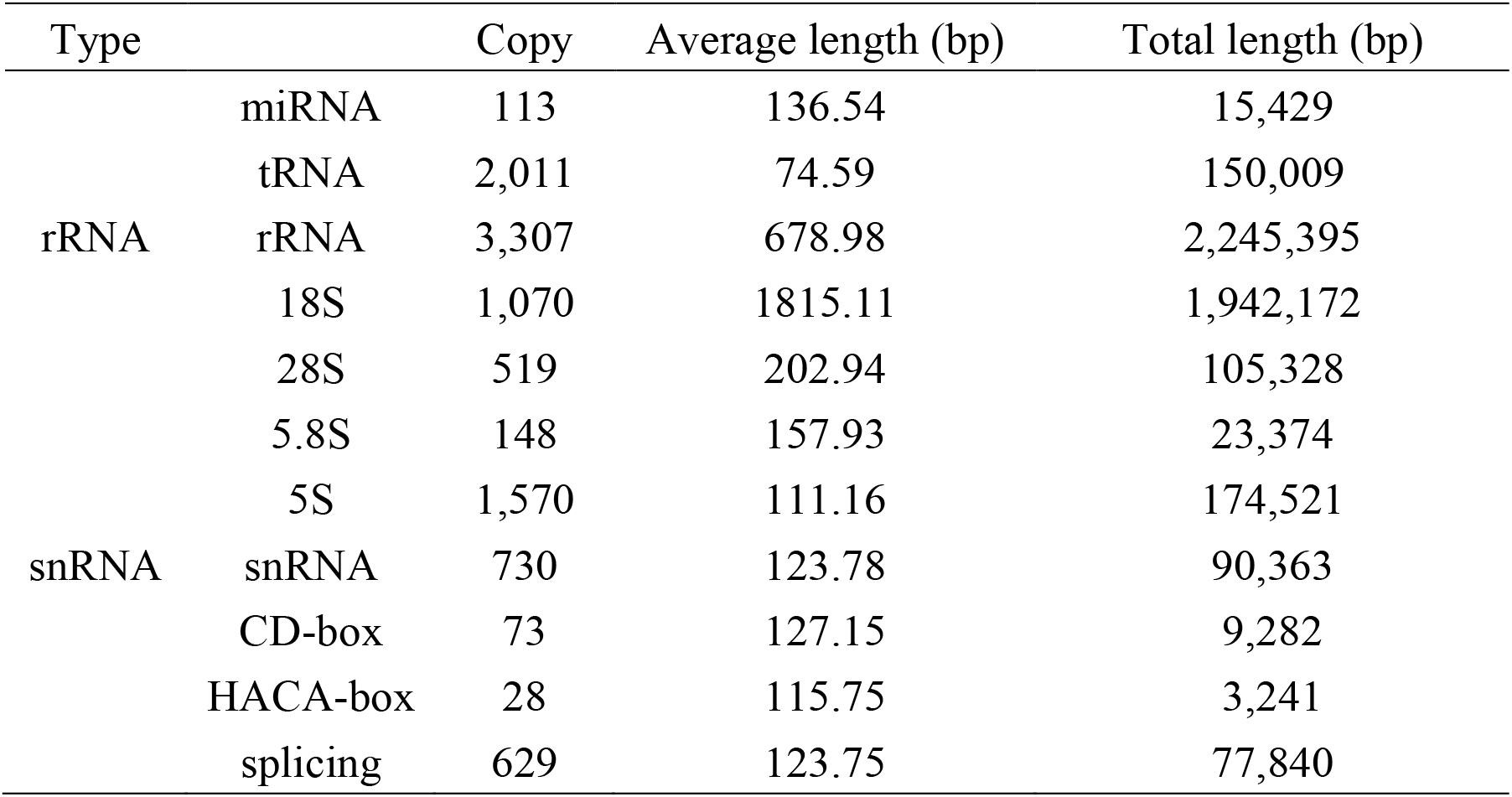
Statistics on the annotation of non-coding RNA of the *P. massoniana* ‘Minlin WY36’genome.

The phylogenetic relationships among the major lineages of existing gymnosperms, particularly Ginkgo and Gnetales, remain a subject of ongoing debate (Stull *et al*., 2021). On the basis of the 46 low-copy gene families derived from ten gymnosperms, five angiosperms, and five seedless vascular plants, we constructed phylogenetic tree (**Figure 1C**). The results support Gnetales as a basal lineage within gymnosperms, diverging from other gymnosperms approximately 249.7 million years ago (Mya). *Ginko biloba* and *Cycas panzhihuaensis* form a sister group to conifers, with these two major evolutionary branches diverging around 177.3 Mya. *Taxus wallichiana* **–** *Cunninghamia lanceolata* diverged from Pinaceae approximately 160.4 Mya. Approximately 51 Mya, Pinaceae rapidly diversified into other species, and *P. massoniana* ‘Minlin WY36’ diverged around 12.6 Mya.

A comparative analysis of orthologous genes among *P. massoniana* ‘Minlin WY36’ and the genomes of 19 species revealed that 1,011 gene families (including 3,074 genes) were unique to the *P. massoniana* ‘Minlin WY36’ genome. Enrichment analyses found that these genes are especially enriched in the Kyoto Encyclopedia of Genes and Genomes (KEGG) terms of ‘plant-pathogen interaction’, ‘Flavonoid biosynthesis’, ‘isoflavanoid biosynthesis’, ‘Fatty acid biosynthesis’, and ‘Fatty acid degradation’, ‘Fatty acid elongation’ (**Figure 1D**). Analysis of orthologous gene family expansion and contraction indicated 2,022 expanded gene families in the *P. massoniana* ‘Minlin WY36’ genome, of which 370 (including 3,868 genes) exhibited significant expansion (*P*<0.01). Additionally, 467 gene families were contracted, with 108 gene families (including 663 genes) showing a significant contraction (**Figure 1C**). Enrichment analyses revealed that the significantly expanded gene families were enriched the KEGG pathways related to ‘terpenoid backbone biosynthesis’, ‘diterpenoid biosynthesis’, ‘flavone and flavonol biosynthesis’, and ‘plant-pathogen interaction’, ‘fatty acid degradation’ (**Figure 1E**). Resin is a major metabolite of pine plants and serves as a physical and chemical defense barrier against insect and pathogen attacks (Liu *et al*., 2021). Resin mainly consist of terpenoids, flavonoids, lignin-like compounds, and phenolic compounds, among which terpenoids are the predominant components (Li *et al*., 2013). The genes that are specific and significantly expanded in *P. massoniana* ‘Minlin WY36’ are enriched in pathways related to plant-pathogen interactions, terpenoid metabolism, and flavonoid metabolism.

## Methods

### DNA extraction and sequencing

All genome-sequencing materials used in this study were collected from adult *P. massoniana* ‘Minlin WY36’ grown in Youxi County, Fujian Province, China. Total genomic DNA was extracted from young leaves using the cetyltrimethylammonium bromide method (CTAB) for Illumina and PacBio sequencing. We used the PacBio Sequel II platform to perform 26-cell single-molecule real-time DNA sequencing.

### Genome size estimate

Before genome assembly, filtered Illumina reads were used to estimate the genome size, heterozygosity, and repeat content of *P. massoniana* ‘Minlin WY36’. Then, a genome survey was performed through *K*-mer analysis using Jellyfish v2.1.4 (Marçais and Kingsford, 2011) and GenomeScope (Vurture *et al*., 2017).

### Genome assembly and Hi-C scaffolding

Assemble *P. massoniana* ‘Minlin WY36’ genome using PacBio data. Canu v1.9 (https://github.com/marbl/canu) (Koren *et al*., 2017) was used to correct the error of the PacBio clean data. We assembled the corrected data using Smartdenovo v1.0 (Ruan, 2016). The Contig-level assembly results were aligned to the genome using Juicer (Kent, 2002), and the alignment results were filtered to remove incorrectly aligned reads. Next, 3D-DNA (Ruan, 2016) was used to perform preliminary clustering, ordering, and orientation of the genomic sequences. To prevent excessive fragmentation of the genomic sequences, we manually adjusted the preliminary clustering results from 3D-DNA using visualization software Juicebox (https://github.com/aidenlab/Juicebox). This manual adjustment involved removing redundancies, repositioning, and re-clustering, as necessary. The final pseudochromosomes were manually constructed and adjusted. To evaluate the results of the Hi-C assembly, we constructed an interaction heat map of Hi-C assembly chromosome.

### Gene prediction and annotation

Annotation information from five closely related species (*Amborella trichopoda, G. biloba, G. monatum, P. abies*, and *P. taeda*) was used for homology-based predictions. From the structurally intact genes obtained through homology-based prediction, we randomly selected 5,000 genes to train the de novo models of the target species using the ab initio prediction software, Augustus and SNAP (Johnson *et al*., 2008). Simultaneously, the RNA-seq data were aligned to the genome using HISAT v2.1.0 (Kim *et al*., 2015) and the resulting transcripts were assembled using StringTie v1.3.4d (Pertea *et al*., 2015). Finally, Maker v2.31.8 (Holt and Yandell, 2011) software was used to integrate homology-based predictions. In addition, the gene sets obtained from gene structure annotation were aligned with known protein databases and other databases such as SwissProt (Boeckmann, 2005) (http://www.uniprot.org/), TrEMBL (http://www.uniprot.org/) (Boeckmann, 2005), KEGG (Kanehisa and Goto, 2000) (http://www.genome.jp/kegg/), and InterPro (https://www.ebi.ac.uk/interpro/) (Zdobnov and Apweiler, 2001) and were aligned using Blast v2.2.26 (Altschul *et al*., 1990).

### Identification of non-coding RNA, and repetitive sequences

Non-coding RNAs include RNA with a variety of known functions, such as microRNAs (miRNAs), ribosomal RNA (rRNA), and transfer RNA (tRNA). Based on the structural characteristics of tRNA, software tRNAscan-SE 1.3.1 (http://lowelab.ucsc.edu/tRNAscan-SE/) (Lowe and Eddy, 1997) was used to search for tRNA sequences in the genome. Owing to the high conservation of rRNA, rRNA sequences from closely related species was selected as reference sequences, and BLASTN alignment was used to search for rRNA sequences in the genome. Additionally, covariance models from the Rfam family (Sam *et al*., 2005), along with the INFERNAL software (http://infernal.janelia.org/) provided by Rfam, can be used to predict miRNA and snRNA sequence information in the genome.

The process involved identifying sequences similar to known repetitive sequences based on the RepBase v21.12 (Jurka *et al*., 2005) database (http://www.girinst.org/repbase) using RepeatMasker v4.0.7 (http://www.repeatmasker.org) and RepeatProteinMask v4.0.7 (http://www.repeatmasker.org). *De novo* prediction, on the other hand, is based on self-sequence alignment, where a *de novo* (Price *et al*., 2005) repetitive sequence library was first established using RepeatModeler (http://www.repeatmasker.org) and LTR_FINDER v1.06 (http://tlife.fudan.edu.cn/ltr_) (Benson, 1997). This library was then used for prediction using the RepeatMasker software. Additionally, Tandem Repeats Finder v4.09 (http://tandem.bu.edu/trf/trf.html) (Price *et al*., 2005) was used to search for tandem repeats within the genome.

### Ortholog detection

Single/low-copy and multicopy gene families can be obtained by identifying orthologous genes and cluster analysis of gene families. A protein dataset that included ten gymnosperms, five angiosperms, and five seedless vascular plants was constructed. Pairwise BLASTP (Altschul *et al*., 1990) alignments were performed on the protein dataset, followed by a clustering analysis using OrthoFinder (Emms and Kelly, 2015).

### Phylogenetic reconstruction

We constructed a species phylogenetic tree using low-copy gene families from *P. massoniana* ‘Minlin WY36’ and 19 other plants and identified 46 low-copy families among the 20 species. Then, the dataset used for phylogenetic analysis was employed in the MCMCTree program within PAML4.9 (Ziheng, 2007) to estimate the divergence times. The nucleotide substitution model was set as the GTR model and the molecular clock model was set as the independent rate model. The MCMC process included 500,000 burn-in iterations and 1,500,000 sampling iterations, with sampling conducted every 150 iterations. The same parameters were run twice to obtain stable results. The phylogeny was calibrated using various fossil records or molecular divergence estimates by placing soft bounds on the split nodes of *G. montanum* – *G. biloba* (230–282 Ma), *Arabidopsis thaliana* – *G. montanum* (289–330 Ma), *Azolla pinnata* – *A. thaliana* (392–422 Ma), and *Physcomitrella patens* – *A. thaliana* (465–533 Ma).

### Gene family expansion and contraction

For the well-clustered gene families and the constructed phylogenetic tree structures, we performed gene family contraction and expansion analysis on the selected 20 species using CAFÉ 4 (De Bie *et al*., 2006). We conducted KEGG and GO enrichment analyses focusing on genes belonging to the notably expanded gene family within the *P. massoniana* ‘Minlin WY36’ genome.

## Acknowledgments

This project was supported by the Forestry Peak Discipline Construction Project of Fujian Agriculture and Forestry University (72202200205), the National Key Research and Development Program of China (No. 2203YFD1600504), and Fujian Forestry Science and Technology Program (No.2020-9).

## Competing interests

The authors declare no competing interests.

## Notes

### Competing Interest Statement

The authors have declared no competing interest.

### Summary of Updates

Update the Acknowledgments to show the project support received for this research: This project was supported by the Forestry Peak Discipline Construction Project of Fujian Agriculture and Forestry University (72202200205), the National Key Research and Development Program of China (No. 2203YFD1600504), and Fujian Forestry Science and Technology Program (No.2020-9). And competing interests: The authors declare no competing interests.

## Reference

Altschul, S. F. et al. Basic local alignment search tool. Journal of Molecular Biology 215(3): 403– 410 (1990).

Benson, G. Tandem repeats finder: a program to analyze DNA sequences. Nucleic Acids Research 27(2): p. 573–580 (1997).

Boeckmann, B. et al. The SWISS-PROT protein knowledgebase and its supplement TrEMBL in 2003. Nucleic Acids Research 1(1): p. 365–370 (2003).

De Bie, T. et al. CAFE: a computational tool for the study of gene family evolution. Bioinformatics 22(10): 1269–1271 (2006).

Emms, D. and Kelly, S. OrthoFinder: solving fundamental biases in whole genome comparisons dramatically improves orthogroup inference accuracy. Genome Biology 16: 157 (2015).

Gernandt, D.S. et al. Molecular phylogeny of Pinaceae and Pinus. In IV International Conifer Conference 615:107–114 (1999).

He, Y. et al. Potential geographical distribution and its multi-factor analysis of Pinus massoniana in China based on the maxent model. Ecological Indicators 154: 110790 (2023).

Holt, C. and Yandell, L. MAKER2: an annotation pipeline and genome-database management tool for second-generation genome projects. BMC Bioinformatics 12(1): p. 491 (2011).

Johnson, A.D. et al. SNAP: a web-based tool for identification and annotation of proxy SNPs using HapMap. Bioinformatics 24(24): 2938–2939 (2008).

Jurka, J. et al. Repbase Update, a database of eukaryotic repetitive elements. Cytogenet Genome Res. 110(1–4): 462–7 (2005).

Kanehisa, M. and Goto, S. KEGG: kyoto encyclopedia of genes and genomes. Nucleic Acids Research 28(1): p. 27–30 (2000).

Kent, W. J. BLAT-the BLAST-like alignment tool. Genome Research 12, 656–664 (2002).

Kim, D. et al. HISAT: a fast spliced aligner with low memory requirements. Nature Methods 12(4): 357 (2015).

Koren, S. et al. Canu: scalable and accurate long-read assembly via adaptive k-mer weighting and repeat separation. Genome Research 27: 722–736 (2017).

Li, B. et al. Chemical constituents and biological activities of Pinus species. Chemistry & Biodiversity 10(12): 2133–2160 (2013)

Liu, B. et al. Two terpene synthases in resistant Pinus massoniana contribute to defence against Bursaphelenchus xylophilus. Plant Cell Environ. 44: 257–274 (2021).

Lowe, T. M. and Eddy, S. R. tRNAscan-SE: a program for improved detection of transfer RNA genes in genomic sequence. Nucleic Acids Research 25(5): 955–964 (1997).

Marçais, G. and Kingsford, C. A fast, lock-free approach for efficient parallel counting of occurrences of k-mers. Bioinformatics 27(6): 764–770 (2011).

Nystedt, B. et al. The Norway spruce genome sequence and conifer genome evolution. Nature 497: 579–584 (2013).

Peng, S. L. et al. Genetic diversity of Pinus massoniana revealed by RAPD markers. Silvae Genetica 52(2): 60–63 (2003).

Pertea, M. et al. StringTie enables improved reconstruction of a transcriptome from RNA-seq reads. Nature Biotechnology 33(3): 290 (2015).

Price, A. L. et al. De novo identification of repeat families in large genomes. Bioinformatics 21 (suppl_1): p. i351 (2005).

Stamatakis A. RAxML Version 8: A tool for phylogenetic analysis and post analysis of large phylogenies. Bioinformatics 30 (9): 1312–1313. 6 (2014).

Ruan, J. Ultra-fast de novo assembler using long noisy reads. Available at https://github.com/ruanjue/smartdenovo (2016).

Sam, G. J. et al. Rfam: annotating non-coding RNAs in complete genomes. Nucleic Acids Res 33, 121–124 (2005).

Stull, G.W. et al. Gene duplications and phylogenomic conflict underlie major pulses of phenotypic evolution in gymnosperms. Nature Plants 7: 1015–1025 (2021).

Vurture, G. W. et al. GenomeScope: fast reference-free genome profiling from short reads. Bioinformatics 33, 2202–2204 (2017).

Zdobnov, E. and Apweiler, R. InterProScan--an integration platform for the signature-recognition methods in InterPro. Bioinformatics 17(9): p. 847–848 (2001).

Zhang, Y. et al. Divergence among Masson pine parents revealed by geographical origins and SSR markers and their relationships with progeny performance. New Forests 44(3): 341–355 (2013).

Ziheng, Y. PAML 4: phylogenetic analysis by Maximum Likelihood. Molecular Biology and Evolution 24(8):1586–1591 (2007).

